# Spike-based symbolic computations on bit strings and numbers

**DOI:** 10.1101/2021.07.14.452347

**Authors:** Ceca Kraišniković, Wolfgang Maass, Robert Legenstein

## Abstract

The brain uses recurrent spiking neural networks for higher cognitive functions such as symbolic computations, in particular, mathematical computations. We review the current state of research on spike-based symbolic computations of this type. In addition, we present new results which show that surprisingly small spiking neural networks can perform symbolic computations on bit sequences and numbers and even learn such computations using a biologically plausible learning rule. The resulting networks operate in a rather low firing rate regime, where they could not simply emulate artificial neural networks by encoding continuous values through firing rates. Thus, we propose here a new paradigm for symbolic computation in neural networks that provides concrete hypotheses about the organization of symbolic computations in the brain. The employed spike-based network models are the basis for drastically more energy-efficient computer hardware – neuromorphic hardware. Hence, our results can be seen as creating a bridge from symbolic artificial intelligence to energy-efficient implementation in spike-based neuromorphic hardware.

## 1 Introduction

Spiking neural networks (SNNs) constitute the third generation of neural network models (Maass, 1997). The most salient difference to standard artificial neural networks (ANNs) that are usually employed in deep learning is the means of communication between neurons in the network. While ANNs communicate real numbers in each computational step, spiking neurons in SNNs communicate events in an asynchronous manner. The output of a spiking neuron is zero most of the time, only occasionally turning to “1”, which is referred to as spiking or sending a spike. This communication scheme is much closer to the way neurons communicate in the brain and it has some advantages that we will discuss below. In this chapter, we discuss symbolic computations with SNNs. In particular, since we concentrate on tasks that involve temporally sequential input which necessitates recurrent connections in order to combine temporally dispersed information, we consider here networks of recurrently connected spiking neurons (RSNNs). This also reflects the fact that neuronal networks in the brain are highly recurrent. We use the term RSNNs whenever we want to empahsize the recurrent architecture of the considered network. We review the current state of this research direction and in addition present new results which show that surprisingly small SNNs can perform symbolic computations on bit sequences and numbers.

Why should we bother about RSNNs when thinking about neuro-symbolic computation? There are at least two answers to this question. First, the brain is very likely a computing system that merges sub-symbolic connectionist computation with symbolic components. It has been argued many times that a pure subsymbolic system cannot achieve the generalization abilities of symbolic systems where variables can be used in various contexts (Marcus, 2001, 2018; Marcus et al., 2014). Hence, the superb generalization capabilities of the brain likely result from symbolic computations. On the other hand, judged purely from the hardware, it is undisputed that the brain is a neural system. However, if we want to learn from the brain, we should take its basic computational units seriously. The event-based nature of spiking communication in the brain is in many aspects different from communication in ANNs with real-valued neurons. One can in principle quite easily emulate every ANN with an SNN since each real-valued output can be encoded by a spike rate, and several such and more advanced conversion methods exist (Rueckauer and Liu, 2018; Kim et al., 2019). On the other hand, due to the asynchronous and spike-timing based communication in SNNs, the emulation of an SNN with an ANN is much harder (Maass, 1997). Hence, it might not be straightforward to map the solution that evolution has found for symbolic computation to ANNs. Second, the brain is a network of a gigantic number of neurons, and the experience with deep learning has shown that bigger is usually better. Hence, we can expect that neuro-symbolic systems that approach at least some aspects of human intelligence will be huge. To make them practically useful, one should consider energy-efficient hardware solutions. One reason why the brain is extremely energy-efficient is its spike-based communication, where spikes are transmitted only when necessary. Energy-efficient neuromorphic hardware is based on this basic principle utilizing RSNNs (Maass, 2016; Davies et al., 2021; Furber, 2016).

Another lesson we learned from recent artificial intelligence (AI) research is that learning-based approaches are the most promising path to AI systems. Hence, we should not try to construct symbolic systems but rather equip neural networks with inductive biases and then train them to perform symbolic reasoning tasks. Since SNNs can exploit precise spike timing, some functions can be implemented with less resources by SNNs as compared to ANNs (Maass, 1997). In practice however, training of SNNs was problematic as the standard training algorithms for ANNs — Backprop for feed-forward networks and Backprop-through-time (BPTT) for recurrent networks — were not applicable due to the non-differentiability of the spiking output of the neurons. Recently however, several solutions to this problem have been proposed (Bellec et al., 2018, 2020) and it turned out that SNNs can also compete with ANNs in practice. Therefore, there is no longer a reason to resort to ANNs when studying neuro-symbolic computation. Here, we will consider a training algorithm for RSNNs called e-prop (Bellec et al., 2020). This algorithm does not only produce competitive results for RSNNs, it is also biologically plausible and can — as opposed to BPTT — easily be implemented in neuromorphic hardware.

In the following section, we will briefly discuss the basic principles of spike-based computations in the brain and introduce spiking neural network models. We will then touch on the main theories of symbolic computation in the brain. In the main part of this chapter, we will give an overview of the state-of-the-art of neuro-symbolic computation with RSNNs. We will show in particular, through a few examples, that biologically realistic RSNNs can perform symbolic tasks on bit strings and numbers. These results show that tasks in which symbols are manipulated can be learned by models of spiking neural networks and even trained using the biologically plausible learning rule e-prop (Bellec et al., 2020). These results are not only of interest from a biological perspective to understand the mechanisms used by our brain, but also as a computing paradigm for energy-efficient neuromorphic hardware. In that light, our results suggest bridging symbolic artificial intelligence to energy-efficient implementations in spike-based neuromorphic hardware.

## 2 Neural networks of the brain and spiking neural networks

Neurons are the basic computational units of the brain. They are cells specialized for information processing and information transmission via brief (approximately 1 ms) stereotyped electrical pulses, so-called action potentials or spikes. A spike — a short, sudden increase in voltage — has an amplitude of about 100 mV and lasts 1 − 2 ms (Gerstner and Kistler, 2002). Since the spikes of a neuron are all alike, a spike itself (the form of the pulse) does not convey much information. Instead, information can be carried by the number of spikes in a certain time window, the precise timing of a spike in relation to the spikes of other neurons, and the temporal spike pattern. As the shape of the action potential is considered irrelevant, one can abstract over it and define the output of a neuron as a sequence of time points *t*^(1)^*, t*^(2)^*, t*^(3)^*,…* where *t*^(*i*)^ is the *i*-th spike of the neuron. Such a sequence of spike times from one neuron is also called a spike train.

The brain consist of large networks of such neurons, connected via synapses. Incoming spikes to a neuron initiate changes in the membrane potential of the cell body (soma), and occasionally, if a certain voltage threshold is exceeded, the neuron generates a spike that is transmitted via synapses to other neurons.

This process is formalized mathematically in spiking neuron models. These models come at various granularities and abstraction levels, with the common characteristic that their outputs are spike trains.

### 2.1 The leaky integrate and fire (LIF) neuron model

One of the simplest spiking neuron models is the leaky integrate and fire (LIF) neuron model, see Figure 1 (left).

**Figure 1:**
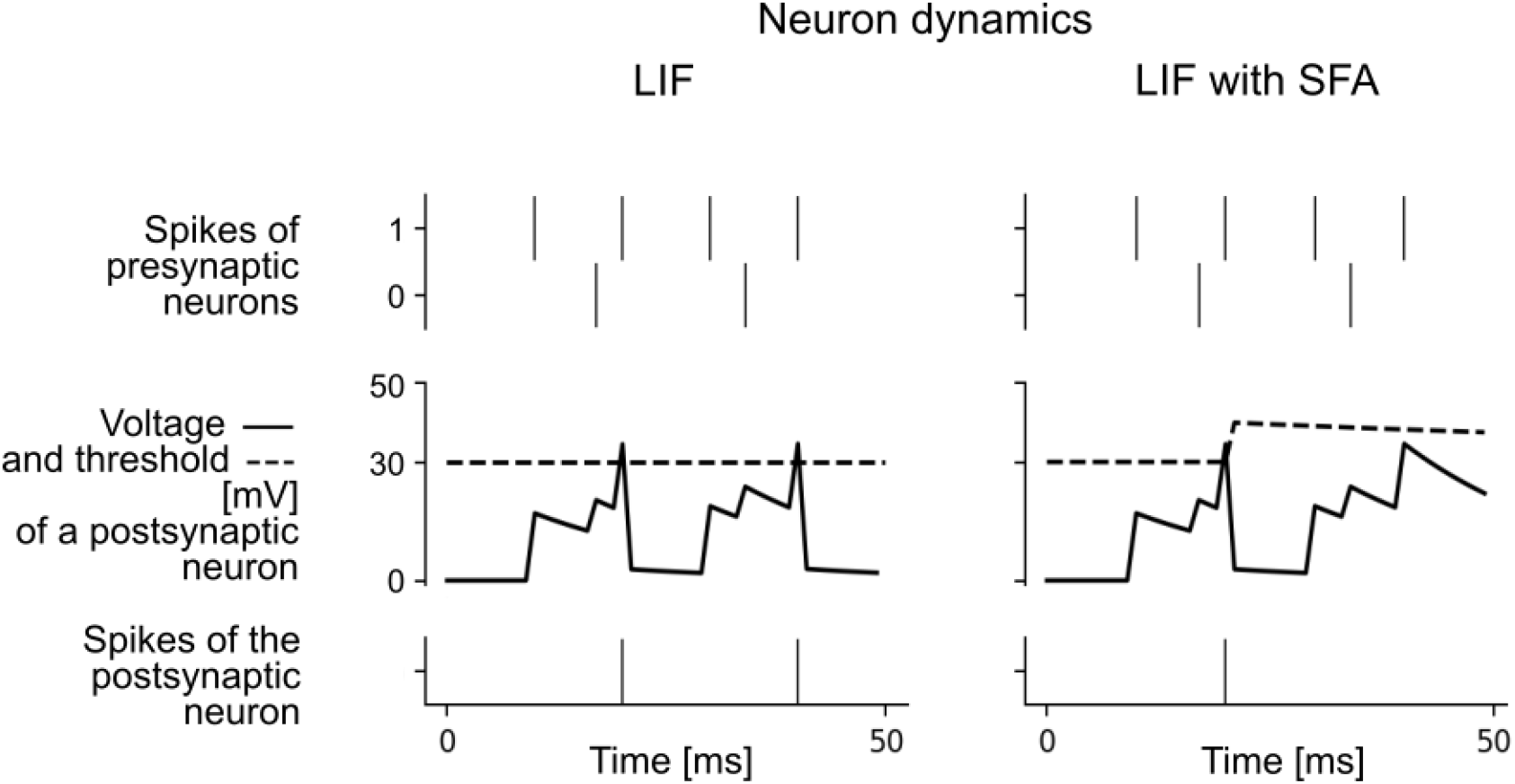
Neuron dynamics for LIF neuron without (left) and with SFA mechanism (right). Top to bottom: The same spike trains of two presynaptic neurons that were connected to a postsynaptic neuron, voltage traces (solid line) and threshold (dashed line) for the postsynaptic neuron without (left), and with spike frequency adaptation (SFA) mechanism (right), output spikes of the postsynaptic neuron. Due to the SFA, the postsynaptic neuron in the right panel outputs only one spike. Neuron parameters used: *τ*_*m*_ = 20 ms, threshold *v*_th_ = 30 mV, *β* = 1.7 mV, *τ*_*a*_ = 100 ms.

It captures the essential temporal dynamics of biological neurons, while abstracting over many — potentially important — aspects of its biological counterparts. The dynamics of the membrane potential *V*_*j*_ of a LIF neuron *j* is described by a first order linear differential equation:

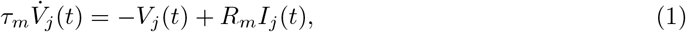

where *τ*_*m*_ is the membrane time constant (on the order of 10 to 30 ms), *R*_*m*_ is the resistance of the cell membrane, and *I*_*j*_ the input current. This equation describes a leaky integrator. The current *I*_*j*_ is integrated over time, however the integration is leaky, i.e., if there is no input current, the membrane potential decays to zero with time constant *τ*_*m*_. The neuron outputs a spike at the time when its membrane potential *V* rises above the threshold *v*_*th*_. After a spike is generated, the neuron dynamics is reset: the threshold value *v*_*th*_ is subtracted from the neuron’s membrane potential, and the neuron enters a refractory period. During the refractory period, the neuron cannot spike.

We consider networks of such neurons where network neurons 1,…, *N* are recurrently connected. We term these networks recurrent spiking neural networks (RSNNs). RSNNs reflect two important biological facts: first, that neurons in the brain are heavily recurrently connected (in contrast to the often used feed-forward architectures in artificial neural networks), and second, that these neurons communicate with spikes. The input current *I*_*j*_(*t*) to neuron *j* in the network is defined as the weighted sum of spikes from external inputs *x* and other neurons in the network *z*:

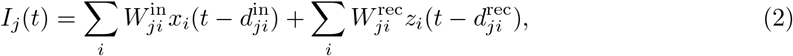

where 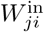 and 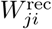 denote the input and the recurrent synaptic weights respectively, and 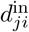 and 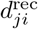 the corresponding synaptic delays. Here, the weights model the efficacies of biological synapses and are similar to the connection weights in standard artificial neural networks. The delay *d*_*ji*_ models the time that passes between an action potential of the presynaptic neuron *i* and the voltage response at the soma of the postsynaptic neuron *j*.

Neurons are usually simulated in discrete time. Writing (1) in discrete time with time-step *δt* and incorporating the reset after a spike, we obtain

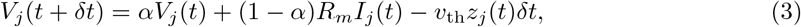

where 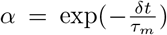 and 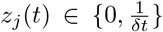 represents the spike output of neuron *j*. The term –*v*_th_*z*_*j*_ (*t*)*δt* implements the voltage reset after each output spike. The output *z*_*j*_ of the neuron is obtained by comparison with a voltage threshold *v*_th_

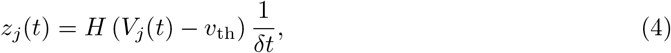

with *H*(*x*) = 0 if *x* < 0 and 1 otherwise being the Heaviside step function.

### 2.2 Enhancing temporal computing capabilities of RSNNs with Spike Frequency Adaptation (SFA)

Upon inspection of the membrane potential dynamics of a LIF neuron, (1), we observe that spiking neurons have some internal memory about previous inputs on the time scale of the membrane time constant *τ*_*m*_, which is on the order of 10’s of milliseconds. This is a difference from standard artificial neurons in deep learning where the output is an instantaneous function of the momentary input. A notable exception are long short-term memory (LSTM) units (Hochreiter and Schmidhuber, 1997), which turned out to be very powerful units in artificial recurrent neural networks. There, so-called memory cells can store their internal state arbitrarily long, which turned out to boost the memory capabilities of recurrent ANNs. The membrane time constant of spiking neurons is however too short to account for short-term memory at behaviorally relevant time scales of seconds in biological organisms. Instead, it has been shown that biological mechanisms at longer time scales can be utilized to boost temporal computing capabilities of RSNNs (Bellec et al., 2018; Salaj et al., 2021). One such mechanism is spike frequency adaptation (SFA), which can easily be incorporated in the standard LIF neuron model.

In contrast to LIF neurons, the threshold of a LIF neuron with SFA is dynamic, see Figure 1 (right). After each spike that the neuron produces, its firing threshold is increased by a fixed amount, and then it decays exponentially back to zero. The effective threshold *A*_*j*_ (*t*) of a LIF neuron *j* with SFA is given by the equations

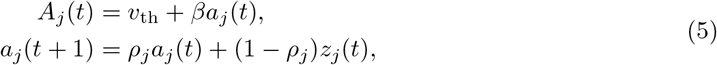

where *v*_th_ is the constant baseline of the firing threshold and *β* > 0 scales the amplitude of the activity-dependent component *a*(*t*). The parameter 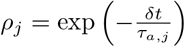 controls the speed by which *a*_*j*_(*t*) decays back to 0, with *τ*_*a,j*_ as the adaptation time constant of neuron *j*. Experimental data show that many neurons in the brain exhibit SFA with time constants *τ*_*a,j*_ from hundreds of milliseconds up to tens of seconds (Pozzorini et al., 2013, 2015).

### 2.3 Training RSNNs with e-prop

Recurrently connected neural networks are trained on temporal sequences of input-output pairs via Backprop-through-time (BPTT) – an algorithm that unfolds the recurrent network in time and operates in two phases. In the first phase, a forward pass is performed, that is, the network outputs are computed, and a cost function that measures the distance between the network’s outputs and desired values is evaluated. In the second phase, by applying the chain rule of calculus, the derivatives of the cost function with respect to the parameters, i.e., synaptic weights, are evaluated for each time step, and the synaptic weights are updated. This approach requires the full history of the network activity to be stored, hence it is quite unlikely that the brain performs such an algorithm for learning. An alternative to the BPTT algorithm that is more biologically realistic is the e-prop learning rule (Bellec et al., 2020). Unlike BPTT, for updating the synaptic weights of the network this learning rule requires only information locally available to each synapse and neuron – an eligibility trace and a learning signal broadcasted across the network. Eligibility traces model preceding activity in neurons and synapses at the molecular level, whereas learning signals model top-down signals (e.g., dopamine, acetylcholine, and neural firing related to the error-related negativity) that target different populations of neurons and govern the synaptic plasticity, i.e., the process of adjusting the strengths of synaptic weights.

More precisely, consider a cost function *E* that should be minimized in the learning process. The gradient 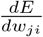 of the cost function with respect to the synaptic weight *w*_*ji*_, needed for the update of this weight can be calculated as follows:

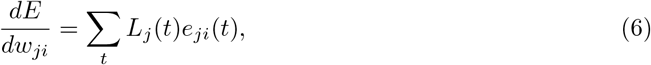

with *L*_*j*_ (*t*) representing the learning signal for neuron *j*, and *e*_*ji*_(*t*) the eligibility trace for the synapse from neuron *i* to neuron *j*. In e-prop, one uses an approximate learning signal

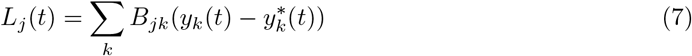

which captures errors occurring at the output neurons *k* at the current time step *t*, that is, deviations of the outputs *y*_*k*_(*t*) from the targets 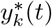 are calculated, and fed back from output neuron *k* to every network neuron *j* via randomly chosen weights *B*_*jk*_. The eligibility trace *e*_*ji*_(*t*) represents a transient memory for the synapse connecting the presynaptic neuron *i* and postsynaptic neuron *j*, that is, it integrates the history of the synapse up to time *t* based on locally available information. For a LIF neuron, it is based on a low-pass filtered version of the presynaptic spike train and the postsynaptic membrane potential.

The learning signal *L*_*j*_(*t*) only takes into account the current time step *t*, however, the eligibility trace *e*_*ji*_(*t*) accounts for the past of the neuron. Since they are readily available in every time step *t* of the forward phase, as soon as the forward phase is completed, the updates of synaptic weights can be performed, without performing the backward phase as required in BPTT. There is no need for unrolling the recurrent network in time for calculating the errors from previous time steps as in BPTT, and as such, e-prop is an online learning method for training models of recurrently connected neurons. For an illustrative, short summary of the e-prop learning rule, see (Manneschi and Vasilaki, 2020).

## 3 Symbolic computation and arithmetics in the brain

### 3.1 Representation of numbers in the brain

There exists experimental evidence for at least two types of neural codes for numbers in the cortex. It has been shown that individual neurons in the lateral intraparietal area (LIP) exhibit a Gaussian tuning for cardinality when macaque monkeys perform a number matching task (Nieder and Miller, 2003). A population of such neurons, each tuned to a different represented value, can collectively encode a number. To make this more precise, we describe here a simple mathematical model for this coding scheme which will also be used in later sections in our simulations. For a represented number *z*, each neuron *i* in such a population spikes with a firing rate of

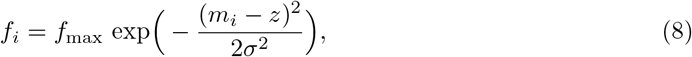

where *f*_max_ is the maximum firing rate of neurons in the population, *m*_*i*_ is the value to which the neuron is tuned to, and *σ* is the standard deviation of the Gaussian tuning curve which is chosen here to be equal for all neurons in the population for simplicity. The preferred values *m*_*i*_ for these neurons are typically chosen equally spaced in the range of numbers that can be represented. Hence, each neuron responds with its maximal firing rate *f*_max_ if its preferred value *m*_*i*_ is represented. When the represented value is off this preferred value, the firing rate decreases according to a Gaussian profile. Such population coding of numeric values between −1 and 1 is illustrated in Fig. 2 for a population of 50 neurons. This coding scheme abstracts over input stimulus properties, as it is preserved over input modalities such as visual or auditory input (Nieder, 2012).

**Figure 2:**
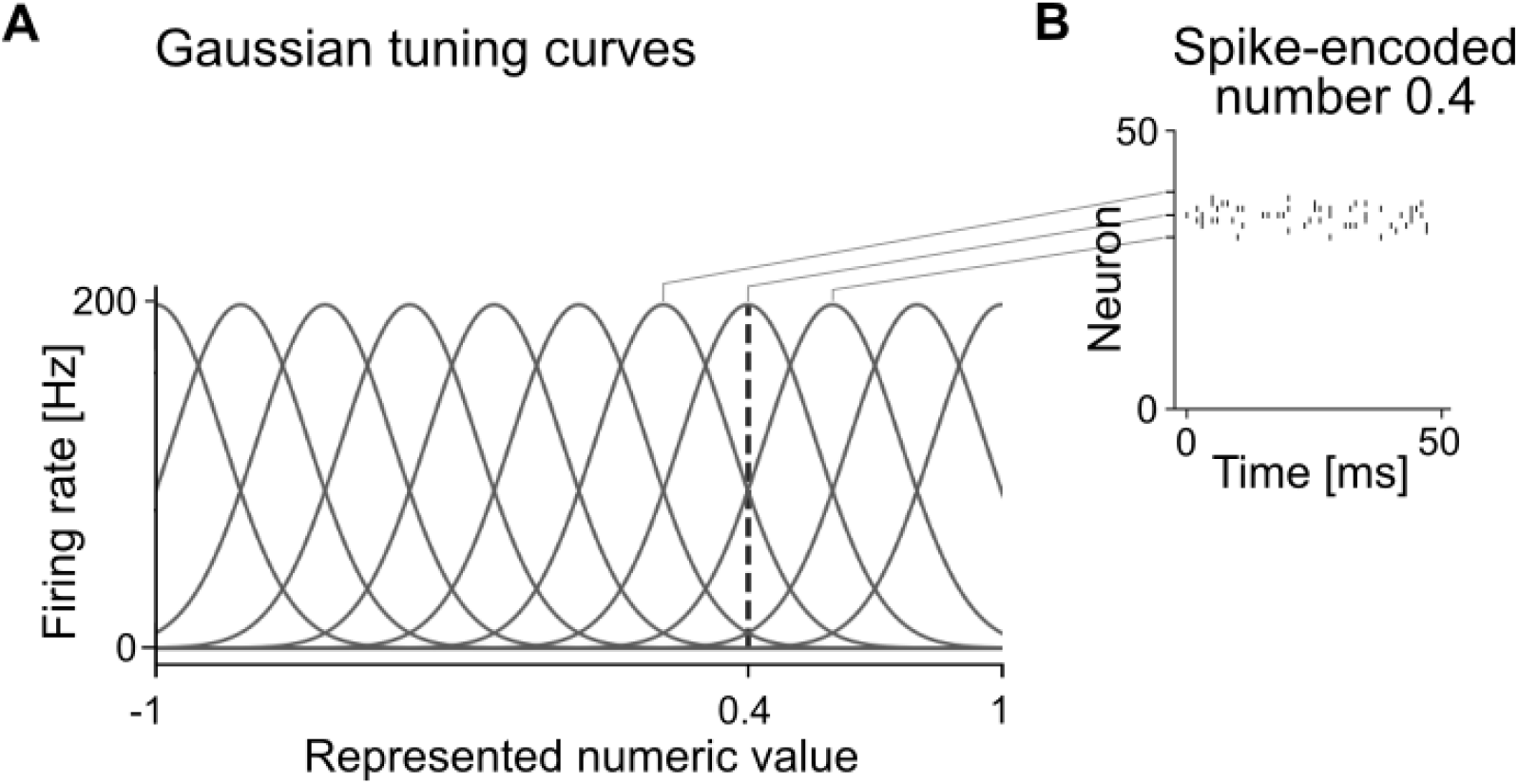
Population coding of numbers. This encoding was used in the middle panel of Figure 4C to encode numbers. (A) The numeric value from [−1, 1] that is represented by a population of 50 neurons is indicated on the x-axis. The y-axis indicates the firing rate of 11 neurons out of this population in response to the corresponding number. Each neuron responds with its maximal firing rate to its preferred value (at the peak of its Gaussian tuning). When the represented value is off this preferred value, the firing rate decreases according to the indicated Gaussian profile. The dashed vertical line indicates one example value of 0.4. The neuron with its tuning curve centered on this value fires at 200 Hz. The two neurons with the shown neighboring tuning curves fire at approximately 80 Hz. (B) Spiking activity of 50 spiking neurons of this population when the number 0.4 was encoded, as indicated in panel A. Each row corresponds to one neuron of the population, where neurons are sorted by their preferred value from −1 to 1. Each tick indicates one spike. Gray lines connect the centers of tuning curves in A with the corresponding neurons in panel B.

In other tasks, some neurons in LIP fire more strongly for larger numbers and others show the inverse behavior, hence exhibiting a spike frequency code (Roitman et al., 2007). It has been hypothesized that these two types of codes may separate the actual entity represented (i.e., the number represented with a Gaussian tuning) from the structure of the number space (a linear firing rate code which captures similarity between numbers) (Summerfield et al., 2020; Behrens et al., 2018). The authors also state that structure-disentangled codes are not unique to numbers and magnitude, rather this might be a general coding principle for structured information (e.g., space). There exists evidence for similar coding principles in humans. EEG activity varies smoothly with numerical distance when Arabic numbers are presented to humans (Spitzer et al., 2017; Teichmann et al., 2018).

### 3.2 Circuits for arithmetic in the brain

Little is known about the specific circuit mechanisms that implement arithmetic operations in the brain. Several cortical regions are activated during arithmetic tasks. Here, the strength of activation of specific areas often depends on the type of task (e.g. approximate versus exact), task difficulty (e.g. 2 + 3 versus 27 × 32), and expert level of the subject (Zamarian et al., 2009). In general, frontal areas are more strongly involved in more complex calculations (Semenza et al., 1997). Interestingly, several lines of research indicate that growing expertise leads to the increased involvement of specific parietal areas whereas frontal areas become less strongly involved (Zamarian et al., 2009). This supports the theory that arithmetic operations can be performed by more general-purpose processing, but practice may train special purpose circuits which can perform the trained tasks with less effort. Our novel results below show that training of complex arithmetic operations can be performed on surprisingly small SNNs.

### 3.3 Symbolic computation

A symbol has meaning and significance; it refers to ideas, objects, relationships and concepts (Gazzaniga et al., 2009). There are a few important aspects to consider when studying the emergence of meaning from symbols, for example, the establishment of links between symbols and objects/concepts that they represent, learning symbolic meaning from context, or generalization over a range of instances of symbols. Also, different brain regions are involved in general meaning processing, but semantic integration mechanisms draw on higher cortical areas of the neocortex, such as prefrontal, posterior parietal, anterior, inferior, and posterior temporal cortex (Pulvermüller, 2013).

Unlike computations performed directly on values such as the activities of sensory receptors, symbolic computation addresses the question of the manipulation of symbols using rules. A symbolic expression can be expressed using variables that are instantiated by concrete values. In algebra, abstract variables *x* or *y* can be substituted by numbers; in language, terms *subject* or *verb* can be replaced by words. The abstract terms act as placeholders, and they are a higher-order representation of knowledge. Through the concept of variable binding — how the placeholders are instantiated by concrete values — one can study how computations performed by neurons give rise to cognitive processes, and eventually elucidate the underlying process in the brain which contribute to language and abstract thought (Marcus et al., 2014).

Symbols can represent categories (e.g., *cat*, *dog*), variables (e.g., *x*, *y*), computational operations (e.g., ‘+’, ‘−’, *concatenate*, *compare*), and individuals (e.g., *Felix*) (Marcus, 2001). The mind is a system that represents variables, operations over variables, and structured representations, capable of distinguishing between categories and individuals. A possible physical implementation of such a system would be a set of buckets, with the bucket’s content representing an instantiation of a category represented by the bucket. Digital computers already use binary registers that correspond to buckets being either empty or full (bits 0 or 1), and the operations over the registers are performed in parallel. Next, each variable should correspond to an anatomically defined region or register. For example, the symbol *“black”* in *“black cat”* or *“black sedan”* should apply equally, irrespective of the categories animals and cars that may be represented in widely separated regions of the cortex (Marcus et al., 2014).

Several neural network models have been proposed for the implementation of symbolic computation via variables and variable-binding mechanisms that bind specific content to a given variable. Pointer-based models (e.g., (Zylberberg et al., 2013)) assume that pointers are implemented by single neurons or co-active populations of neurons which are synaptically linked to content. Another class of models is based on the idea of indirect addressing. These models assume that a neural space for a variable encodes an address to another brain region where the corresponding content is represented (Kriete et al., 2013; O’Reilly, 2006). In anatomical binding models, different brain areas are regarded as distinct placeholders (Hayworth, 2012; Hayworth and Marblestone, 2018). In Dynamically Partionable Autoassociative Neural Networks (DPANN) (Hayworth, 2012), a given symbol corresponds to a global attractor state of the dynamics of a large-scale network that links many regions of the brain. Information flow between particular sets of registers is achieved through turning on and off subsets of synapses, i.e., gating. In neural blackboard architectures, one assumes that besides assemblies for content, there exist also assemblies for structural entities such as phrases, semantic roles, et cetera. Specialized circuits (so-called gating circuits and memory circuits) are then used to bind content to structure (van der Velde and de Kamps, 2006). This way, complex syntactic structures can be built. Vector-symbolic architectures (Plate, 1995) use high-dimensional vectors to represent the symbolic concepts encoded by activity patterns of a group of neurons and a defined set of mathematical operations. Both the variables and potential role filters, such as subject, object or verb, are represented as vectors. Compositional operations typical for symbol manipulation are then implemented through a mathematical transformation of these vectors. Activity patterns for binding, for example, a subject and an instance, are achieved through multiplication of two vectors (Eliasmith et al., 2012). Most of these models rely on specifically constructed circuitry that performs the binding operations. In contrast, a rather generic SNN model was proposed in (Müller et al., 2020). In this assembly pointer model, binding emerges through local Hebbian plasticity processes from the activation of assemblies in areas that contain variables to areas that contain content.

## 4 Prior work on symbolic computations on bit strings and numbers with SNNs

Some of the above mentioned neural network models for variable binding have been implemented in SNNs (e.g., (Eliasmith et al., 2012; Müller et al., 2020)). In this review, we concentrate more specifically on symbolic computations on numbers, and in contrast to the above mentioned models, we propose a more generic approach where SNNs are trained to perform specific tasks.

In a recent work (Salaj et al., 2021), a spiking neural network was trained on two tasks based on a rule that was given at the beginning of each trial. For the same input sequence, the task of the network was either to reproduce the sequence in the same order, if the rule was *duplicate*, or the sequence in reversed order if the rule was *reverse*. The SNN had to learn to observe data stimuli (symbol sequences) given as input, and two different rules on how to manipulate them, also given as an input, independent of each other, then to use both data stimuli and the given rule to produce an output. After training, the SNN was able to produce sequences with all correct symbols in 95.88% of test trials. Through an analysis of spiking activity on the test trials, groups of neurons that had specialized in encoding structural information of the task, such as symbol identity, position, and task identity, were identified. The analysis revealed that many neurons specialized in encoding non-linear combinations of conditions, indicating a non-linear mixed selectivity of neurons. In other words, while some neurons showed some preference in their tuning to specific aspects necessary to solve the tasks, there was no clear-cut encoding of symbols by individual neurons in this trained circuit — different from what one would usually do to construct a neural network to perform these operations. Such mixed selectivity of neurons has also been observed in higher cortical areas (Rigotti et al., 2013).

In another experiment of (Salaj et al., 2021), it was shown that SNNs can learn to follow dynamic rules, i.e., make decisions, for example, choose the “Left” or “Right” button, depending on the currently active rule and specific subsequences of symbols relevant for that particular rule. In a long sequence of symbols, the currently active rule, relevant and distractor symbols appeared sequentially. Even for humans, the task is quite demanding, since it requires not only the maintenance and an occasional update of the active rule, but also maintaining relevant symbols and ignoring irrelevant ones until the target subsequence is complete. Such a task requires a hierarchical working memory – higher-level memory for maintenance of the active rule, and lower-level memory for maintenance of the symbols relevant in the context of the active rule, especially because the distractor symbols occur rather frequently. A generic SNN was able to solve this task almost perfectly without requiring a special-purpose architecture. In 97.79% of trials consisting of sequences of 90 symbols, all choices of the button “Left” or “Right” were correct. This result shows that cognitively demanding sequence processing tasks can be solved by generic SNNs.

## 5 New results on spike-based symbolic computation on bit sequences and numbers

In this section, we present new results on how SNNs can perform symbolic computations on bit sequences and numbers. Human infants can perform mental arithmetic using nonsymbolic, approximate number representations. Even before children learn algorithms for manipulating numerical symbols and formal mathematics, they are able to perform approximate addition and subtraction of large numbers. In such experiments, children performed well above chance – they were able to do so by comparing quantities instead of providing an exact solution (Gilmore et al., 2007). Certainly, we use non-symbolic representations before we master symbolic arithmetic, which takes years (Gilmore et al., 2010). Some authors refer to this capability as “the number sense” (Dehaene, 2011).

There are many open questions: How do humans perform arithmetic? How are the numbers encoded? What are the mental representations behind the comparison of two numbers or quantities? For the case of symbolic arithmetic, we face even more general questions – those of symbol manipulation and sequence processing.

Mastering symbolic representations and symbol processing, in general, helps us perform cognitively demanding tasks. Certainly, the human brain is capable of processing and manipulating symbols, or even sequences of symbols, thereby following the abstract relationships between mental concepts (symbols). These abstract relationships are known as rules, and the brain can process sequences of symbols even when rules dynamically change or when the stimulus is novel (Marcus, 2001).

Given a sequence of items, humans are able to infer the rules that generated the sequence and apply the same rule even on sequences of items that they had never encountered before (Liu et al., 2019). If we revisit the question of numbers, for example, Arabic numbers and their representation, we notice that single-digit numbers are represented by symbols 0 to 9. Multi-digit numbers are represented by sequences of symbols, where the position in the sequence is of particular relevance. Moreover, we learned the meaning of the most significant and least significant digit, and that the length of the sequence representing the number carries another crucial piece of information. To compare two numbers given in a symbolic form, we can first rely on the rule of comparing the lengths of sequences. In case the lengths are the same, we immediately proceed by applying the second rule - comparing the corresponding significant digits, iteratively if necessary, starting from the most significant digits towards the least significant ones, but we stop when we can unambiguously conclude which number is greater, or after we compared the least significant digit. In fact, we can perform such sophisticated sequence processing during the comparison of two symbolic numbers with little effort. We have mastered this processing during years of practice through schooling, and we are applying the learned rules rather automatically. This subjective observation is backed up by experiments which suggest that specialized cortical circuits learn complex arithmetic operations during training (Zamarian et al., 2009), see Section 3.2.

We performed two experiments – both of them show that rather small, generic SNNs can be trained to perform demanding sequence processing tasks. In the first experiment, we considered the number comparison task described above, a task that has often been used to study number processing in cognitive science (Sheahan et al., 2021). For simplicity, we used binary numbers instead of previously described Arabic numbers, since the same sequence processing rules for comparison also apply when comparing numbers in any base. Note that, the rules describing how to compare the numbers were not explicitly given to the network, rather, the network learned to perform the task through training.

In the second experiment, an SNN was trained to perform arithmetic operations (addition, subtraction, and multiplication) on real numbers encoded into spiking activity using population coding with Gaussian tuning, a number encoding that has been observed in monkeys, see Section 3.1. Arithmetic expressions — the instructions indicating the rule to be applied on the data stream, i.e., real numbers — were given to the network in a separate stream, and sequentially. The challenge of this task was to incorporate pieces of information processed earlier, that is, to reuse the result of the computation from the previous step while following new instructions. Without explicitly being trained to do so, the network had also to solve the binding problem – between variables that appeared in the expression and the neurons encoding the number. The network learned successfully to process a sequence of arithmetic expressions, and similar to humans, to perform at a level of expertise.

### 5.1 Spike-based symbolic computation on bit strings

The comparison of two numbers is a task often considered in cognitive science (see e.g., (Sheahan et al., 2021) for a recent example). We wondered if an SNN could learn to perform the comparison task on expressions with numbers encoded as bit strings. We refer now to the numbers represented in a binary base as bit strings rather than sequences, since we use the term sequence to denote the (temporal) processing dimension and that symbols appear sequentially, one after the other.

The schematic network architecture is shown in Fig. 3A. The input to the network were binary strings of width 10 bits representing numbers in the binary base. Another input stream represented the relational operator to be applied, where we used *equals* ‘=’, *less than* ‘<’, and *greater than* ‘>’. Variables *x*_1_, *x*_2_, and the relational operator were presented to the network sequentially as Poisson spike trains. A Poisson spike train is a random sequence of spikes with a given rate (i.e., expected number of spikes per second), similar to spike trains observed in the brain. Mathematically, such a spike train is generated by a Poisson point process. Each presentation of a variable or relational operator lasted for 200 ms, with high firing activity coding for 1 and low firing activity coding for 0, see Fig. 3C, top. Only after the full sequence indicating the first number *x*_1_, the second number *x*_2_, and the relational operator *rel_op* has been shown, does the network have all the pieces of information required to determine whether the provided relation *x*_1_ *rel_op x*_2_ was correct. As the answer, the network was required to output “Yes” or “No”.

**Figure 3:**
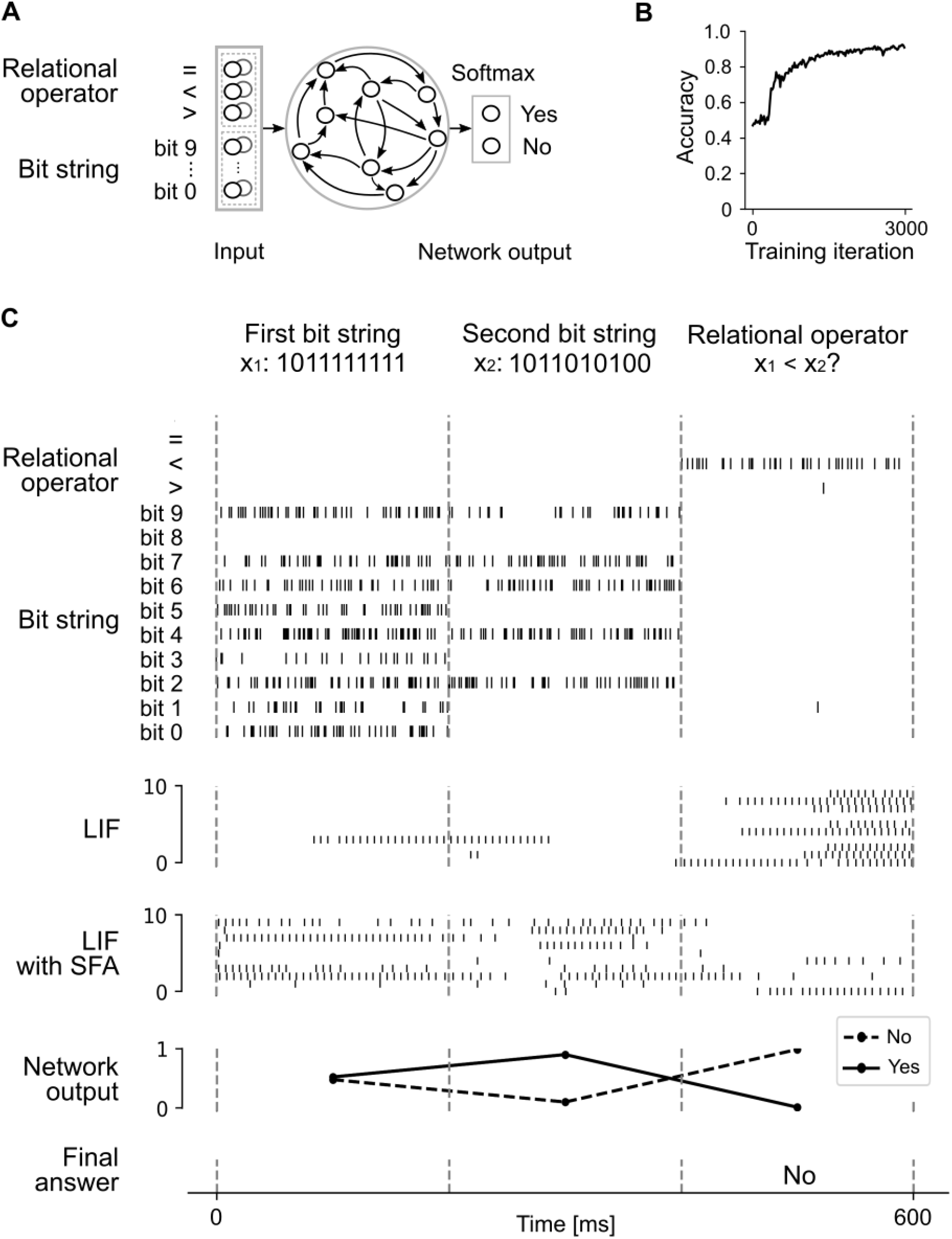
Comparison of two bit strings. (A) The schematic network architecture: input spiking neurons encoding relational operators and bit strings of width 10 (bit 9 the most significant, and bit 0 the least significant bit), recurrently connected LIF neurons, and two output neurons. The network output is the softmax of the ouput neurons producing “Yes/No” answers. (B) Training curve: Accuracy over training iteration. Test accuracy for novel trials was 0.8723. (C) Spiking activity of the network for a trial where the network compared if the first bit string *x*_1_ = 1011111111 was smaller than the second bit string *x*_2_ = 1011010100. Top to bottom: Spiking activity of neurons encoding the relational operator and bit strings (one neuron shown out of 2 for each symbol), 10 sample out of 100 LIF neurons, 10 sample out of 100 LIF neurons with SFA, network output – the softmax applied to readout neurons dedicated to producing the answer, the final (maximum value of those for “Yes” and “No” neurons) and correct answer “No” to the question “*x*_1_ < *x*_2_?”.

We trained a recurrently connected spiking neural network consisting of 200 LIF neurons (100 LIF neurons with SFA, 100 without) on this task using the biologically plausible learning algorithm e-prop (Bellec et al., 2020), for which the accuracy over training iteration is shown in Fig. 3B. The curve was smoothed using bins of 20 values. Fig. 3C illustrates a trial where the network had to answer whether the binary number *x*_1_ = 1011111111 was smaller than binary number *x*_2_ = 1011010100, for which the final answer is “No”. Tested on previously unseen instances *x*_1_ *rel_op x*_2_, that is, combinations of bit strings and relational operators that were not used during training, the proportion of correct answers (accuracy) achieved was 0.8723 ± 0.0235 (mean ± standard deviation), averaged over 5 different network initializations.

This result demonstrates that rather small SNNs can learn to perform arithmetic tasks using a biologically plausible learning algorithm. In this task, numbers were represented in a symbolic manner (i.e., using symbol strings as in Arabic numbers) rather than as magnitudes, and the network had no prior knowledge on the structure of this number encoding or on the structure of the task. Nevertheless, the network was able to generalize to novel task instances, indicating that explicitly designed symbolic computations are not necessary for generalization for this task.

### 5.2 Evaluation of nested arithmetic expressions

One of the most basic arithmetic tasks is the evaluation of an operation applied to two numbers such as 12 + 4. One can replace numbers in such expressions by variables to obtain an expression such as *x*_1_ +*x*_2_. The solution to the expression can be obtained when the values of the variables are defined. For example, defining *x*_1_ = 12 and *x*_2_ = 4, the result for the above expression is 16. This calculation demands a basic type of computation where values need to be assigned to variables which then have to be combined by some arithmetic operation. We wondered whether such basic symbolic computations on numbers and variables could be learned by SNNs. However, we took a further step by considering long nested expressions built based on this basic template with three basic arithmetic operators addition ‘+’, subraction ‘–’, and multiplication ‘*’. An example of such a complex expression is

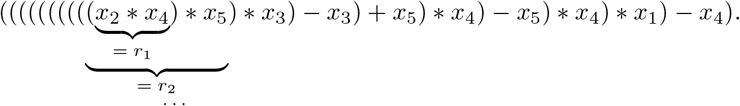

The expressions we considered have a nested structure, where the result *r*_1_ of the evaluation of the expression in the innermost nesting level, here, (*x*_2_ * *x*_4_) is used to evaluate the result *r*_2_ for the next nesting level, and so on, until the final result is obtained. We considered nested expressions of 10 operators combining 5 variables, leading to 9 intermediate results *r*_1_*,…, r*_9_ and the final result *r*_10_ of the evaluation.

More precisely, we define the task as the sequential evaluation of expressions in 10 steps *i* = 1,…, 10 as follows:

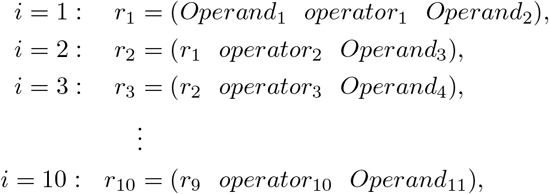

where in the first step (*i* = 1), *Operand*_1_ and *Operand*_2_ are drawn uniformly over {*x*_1_*,…, x*_5_} without replacement, and *operator*_1_ was drawn uniformly over {+, *} (the operator ‘–’ was omitted in this step to avoid the need of encoding the order of *Operand*_1_ and *Operand*_2_). For the following steps, each of *operator*_2_*,…, operator*_10_ was drawn uniformly from {+, –, *}, and each of *Operand*_3_,…, *Operand*_11_ uniformly over {*x*_1_,…, *x*_5_}. This describes the generation procedure for the symbolic part of the task, but numeric values are also required in order to evaluate such nested expressions. We allowed that the values of variables can change after each step *i*. This was done by drawing in each step *i* for each of the variables *x*_1_,…, *x*_5_ a random value from the uniform distribution over [− 1, 1]. An example value assignment and target results for the expression given above are shown in Table 1.

**Table 1:** Evaluation of nested arithmetic expressions with an SNN. One example input sequence with network outputs, targets, and errors. In each input step *i* — each step lasting for 50 ms — an arithmetic expression and numeric values were given to the network, for which the network produced the output *r*_*i*_. Output values were very close to target values, that is, the absolute error was close to zero.

| Step $i$ | Arith. expression | Numeric expression | Output $r_i$ | Target | Absolute error |
| --- | --- | --- | --- | --- | --- |
| 1. | $x_2 * x_4$ | $0.4966 * (-0.1113)$ | -0.0794 | -0.0553 | 0.0241 |
| 2. | $r_1 * x_5$ | $(-0.0794) * (-0.7100)$ | 0.0510 | 0.0564 | 0.0054 |
| 3. | $r_2 * x_3$ | $0.0510 * 0.8209$ | 0.0629 | 0.0419 | 0.0210 |
| 4. | $r_3 - x_3$ | $0.0629 - 0.3314$ | -0.2308 | -0.2685 | 0.0377 |
| 5. | $r_4 + x_5$ | $(-0.2308) + 0.4717$ | 0.2096 | 0.2409 | 0.0313 |
| 6. | $r_5 * x_4$ | $0.2096 * 0.7014$ | 0.1304 | 0.1470 | 0.0166 |
| 7. | $r_6 - x_5$ | $0.1304 - 0.6650$ | -0.5417 | -0.5346 | 0.0071 |
| 8. | $r_7 * x_4$ | $(-0.5417) * (-0.0165)$ | -0.0222 | 0.0089 | 0.0311 |
| 9. | $r_8 * x_1$ | $(-0.0222) * (-0.8182)$ | -0.0128 | 0.0182 | 0.0310 |
| 10. | $r_9 - x_4$ | $(-0.0128) - (-0.4727)$ | 0.4739 | 0.4599 | 0.0140 |

We show here that an SNN can be trained to evaluate such complex arithmetic expressions. The nested expression was given sequentially, starting from the expression in the innermost nesting level (*x*_2_ * *x*_4_ in the above example), together with the numeric assignments to all operands *x*_1_,…, *x*_5_. These inputs were encoded by spiking input populations (see below for the encoding) and presented to the network for 50 ms, see Fig. 4A for a schematic of the input populations. The network was demanded to produce at its readout the intermediate result *r*_1_. Then, the expression for the next nesting level (\**x*_5_ in the above example) was provided for 50 ms and the result *r*_2_ was demanded as the output, and so on, see Fig. 4C for the temporal sequence of inputs to the network.

**Figure 4:**
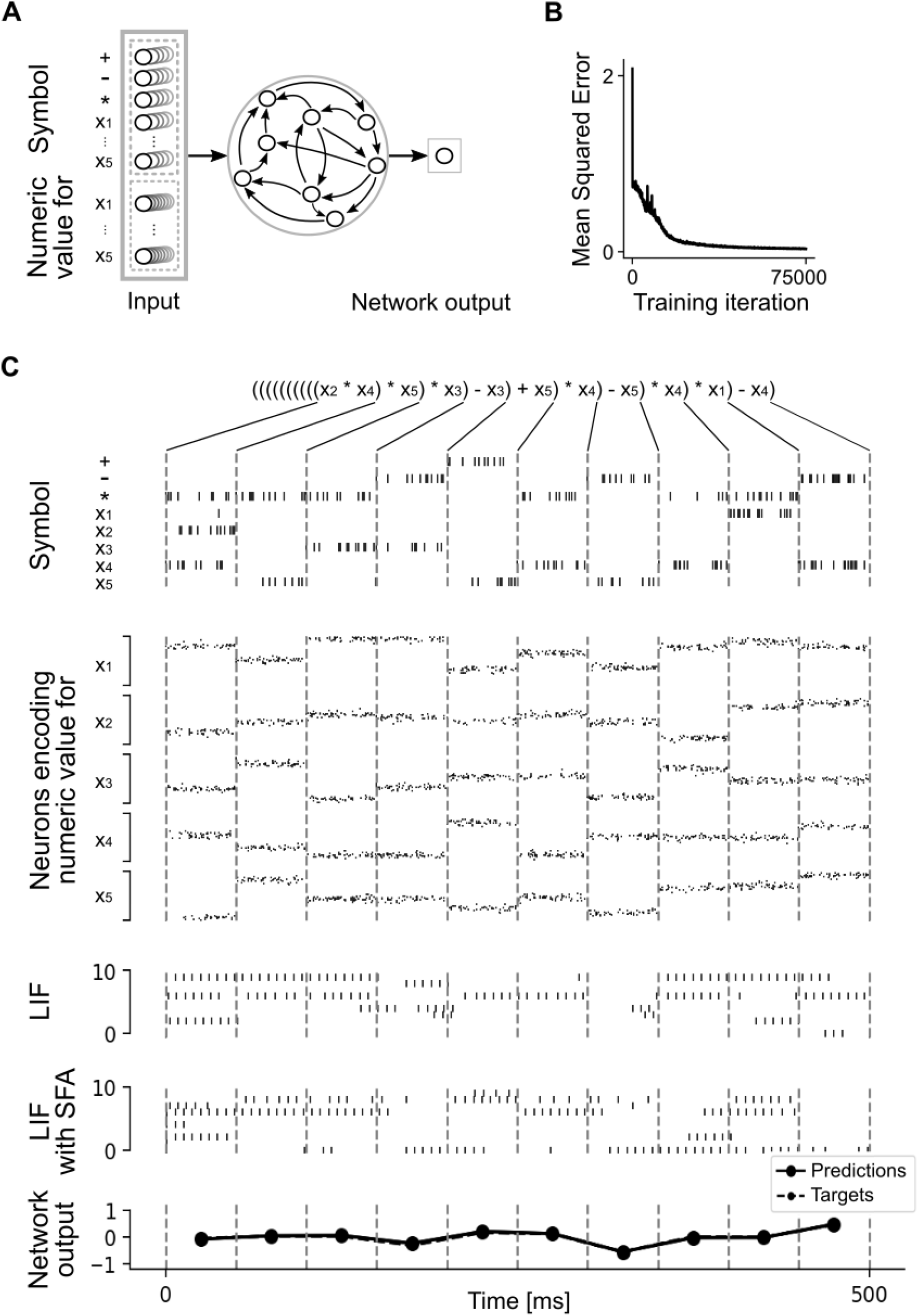
Spike-based arithmetic on numbers and variables. (A) The schematic network architecture consisting of input spiking neurons that encoded symbols used in arithmetic expressions (operators and variables) and numeric values for variables, recurrently connected LIF neurons, and a linear readout neuron that produced the result for each arithmetic expression. (B) Training curve (Mean Squared Error over training iteration) for the network trained to evaluate nested mathematical expressions. The curve was smoothed using bins of 50 values. (C) Spiking activity of the network for the nested arithmetic expression shown at the top. Solid lines from the expression to dashed lines of the spike raster indicate the time when this subexpression was presented to the network. Spike rasters from top to bottom: Spiking activity of neurons encoding the symbols appearing in the arithmetic expression (one neuron shown out of 5 for each symbol), encoding of numeric values that instantiate variables, 10 sample out of 150 LIF neurons (without SFA), 10 sample out of 150 LIF neurons with SFA, real-valued network outputs (predictions) that almost perfectly match targets (and consequently, the dashed line representing targets is difficult to see).

Consequently, the challenges for the network were the processing of symbols with associated numeric values, i.e., it had to learn to bind a concrete value to a variable, and at the same time, carry out the demanded mathematical operations in an online fashion. An example of computations performed by a trained SNN on the previously mentioned nested expression is illustrated in Table 1.

The schematic architecture of a spiking network that we trained is illustrated in Fig. 4A. Symbolic inputs to the network were encoded by subpopulations of neurons producing Poisson spike trains, dedicated to each symbol {+, −, *, *x*_1_,…, *x*_5_} (5 neurons per symbol) and indicating the presence (absence) of a symbol through a high (low) firing rate. Numeric values for each of variables *x*_1_,…, *x*_5_ were encoded by another set of 5 subpopulations of neurons, each consisting of 50 neurons using a population coding scheme. See Fig. 2 for an illustration of population coding of a single number with a population of 50 neurons. Each neuron of this population was assigned a preferred value for which it exhibited its maximal firing rate of 200 Hz. When the represented number was off this preferred value, its rate decreased according to a Gaussian profile. The preferred values of the neurons in this population uniformly tiled the whole space of values between −1 and 1 (Fig. 2A). Hence, for a specific represented value, neurons with preferred tuning close to the value were active, while others remained mostly silent, see Fig. 2B.

As illustrated in Fig. 4A, the spiking activity of the input populations was projected to an RSNN. The output of the network was a single linear non-spiking neuron (a readout neuron), where the numeric value of its output (a real number) was interpreted as the result of the computation. We trained an RSNN consisting of 300 LIF neurons (150 with SFA and 150 without SFA) with all-to-all connectivity using e-prop for 75000 iterations on this task. The objective minimized was the Mean Squared Error (MSE) between the network outputs (predictions) and targets for all intermediate and the final result of the arithmetic expression. The evolution of the network error (MSE) over the training iterations is shown in Fig. 4B. The network achieved an MSE of 0.0341 and a mean absolute error (MAE) of 0.1307 in the mean over all intermediate results, i.e., all computational steps in all of 5000 test trials.

The column “Absolute error” of the example illustrated in Table 1 confirms that the network learned the task very well. The spiking input (encoding of symbols, and of numeric values for variables) and network spiking activity of LIF neurons for the same trial is shown in Fig. 4C, where the bottom panel shows an almost perfect match of outputs (predictions) and targets.

We note that we made sure that the nested symbolic expressions used in the testing trials were different from any nested expression used during training. Also, the used numeric values for the operands were drawn randomly, which, altogether, indicates good generalization capabilities of the trained SNN. This result shows that nested arithmetic expressions can be learned by rather small SNNs to high precision with a biologically plausible learning algorithm. Again, no explicit mechanisms for variable binding were necessary.

## 6 Conclusions

Most studies of computing capabilities of RSNNs have focused on sensory processing tasks. But the brain uses RSNNs also for higher cognitive functions such as symbolic computations, in particular, mathematical computations. We have reviewed recent results, and added two new ones, which show that RSNNs are very good at these tasks. Moreover, they are able to acquire these capabilities through a biologically plausible learning method – e-prop, and they are able to apply learned computing procedures to new instances of the task that never occurred during training. This is somewhat surprising, since RSNNs do not compute in a clocked mode, like digital computers or ANNs. Rather, they compute in an asynchronous mode where also the timing of spikes carries information. Also, the shown examples for such RSNNs operate in a rather low firing rate regime, where they could not simply emulate ANNs by encoding continuous values through firing rates. Thus we arrive here at a new paradigm for symbolic computation in neural networks that provides concrete hypotheses about the organization of symbolic computations in the brain. Furthermore, since RSNNs can be implemented more energy-efficiently in neuromorphic hardware than ANNs, the results that we have discussed also suggest new methods for improving the energy efficiency of computing hardware for symbolic computations.

## 7 Technical details to new results

### 7.1 Network architecture and neuron model

We trained a recurrent spiking neural network of LIF neurons with all-to-all connections. Since in both experiments that we demonstrated temporal integration of pieces of information was crucial for a good performance, a fraction of LIF neurons (one half) was equipped with spike frequency adaptation (SFA). LIF neuron model was described in Section 2.1, and LIF neuron model with SFA in Section 2.2.

The firing threshold of LIF neurons was *v*_th_ = 30 mV. The same value of 30 mV was used as a baseline firing threshold *v*_th_ for LIF neurons with SFA, with an adaptation strength of *β* = 1.7 mV (see (5)). All LIF neurons had a synaptic delay of 1 ms, a refractory period of 5 ms, and a membrane time constant of *τ*_*m*_ =20 ms. The resistance of the cell membrane was *R*_*m*_ = 1 GΩ. A population of input neurons encoded inputs to the recurrently connected neurons in the form of spike trains. The network output was produced by linear readout neurons, from the average activity of recurrent neurons in the network during the time window representing one step of a trial.

### 7.2 Training method e-prop

In all tasks, input, recurrent, and readout weights were trained simultaneously. To train the weights of the network by the learning rule random e-prop (Bellec et al., 2020), we used the Adam optimizer with a learning rate of 0.001. The input and recurrent weights were initialized with values from a standard normal distribution, and then scaled with a factor 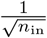, and 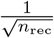, respectively, with *n*_in_ being the total number of input neurons, and *n*_rec_ the number of recurrent neurons in the network. The readout weights (weights from recurrent to output neurons) were initialized with values from a normal distribution with mean zero, and standard deviation 1.05. The unit for all input and recurrent weights was *pAs*.

### 7.3 Tasks

#### Comparison of bit strings

An SNN consisting of 200 recurrently connected neurons was trained for 3000 iterations, with a batch size of 100, to answer the “Yes/No” questions of the form “Is *x*_1_ *Equal to / Smaller than / Greater than x*_2_?”. Inputs to the network were two binary strings *x*_1_ and *x*_2_ of width 10 bits, and in fact, represented binary encoded numbers. Binary strings *x*_1_ and *x*_2_ were given successively in the first and the second step of each trial, then followed by a relational operator =, < or >, given in the third step. In the third, final step, the network had to produce a “Yes/No” answer to the question that got complete, after all three important pieces of information were shown to the network. Each step lasted 200 ms, hence the total duration of the trial was 600 ms. 10-bit binary strings *x*_1_ and *x*_2_ were encoded into spike trains by a population of 20 spiking neurons – two neurons were dedicated to encoding each bit. Relational operators {=, <,>} were encoded using 6 input spiking neurons (2 neurons per operator), in each trial exactly one operator being given in the third step. If a bit represented a value of 1, or if the operator was used in the expression, dedicated neurons fired with a high rate of 200 Hz. To encode a bit value of 0 or that the operator was not used in the expression, neurons fired with a low rate of 2 Hz. Such spike trains encoding the inputs were received by a population of recurrently connected LIF neurons – 100 LIF neurons with SFA and 100 without. Neurons with SFA had an adaptation time constant uniformly distributed from [1, 600]. The linear readout consisted of 2 neurons on which we applied softmax – one neuron was dedicated to “Yes” and one for “No” output. Final answer was determined from the maximum value after the softmax – the higher value indicated either “Yes” or “No”.

The loss function minimized was cross-entropy function with an additional regularization term. The regularization term represented the squared difference between the average firing rate of the recurrent neurons and a target firing rate of 20 Hz, scaled by a factor of 10. On average, in 87.23% of, in total, 10000 testing trials, the network produced the correct answer. The average was calculated over 5 different initializations of networks with the same architecture and trained in the same way (the weights and adaptation time constants of LIF neurons with SFA differed in each run). The inputs in each test trial were novel, i.e., not used during training.

#### Evaluation of nested arithmetic expressions

An SNN consisting of 300 recurrent neurons was trained for 75000 iterations, with a batch size of 50 to perform symbolic computation on nested arithmetic expressions involving numbers and variables. Symbolic input to the network was an arithmetic expression formed by the variables *x*_1_,…, *x*_5_ and arithmetic operators {+, −, *}. In the first step of each trial of length 10, two different variables were drawn randomly to form a valid expression with two operands, and the possible operator was limited to either addition (+) or multiplication (*), to avoid the problem of encoding the order of operands appearing in the expression, which a noncommutative operator subtraction (−) would require. In all other steps of a trial, a variable and an operator were drawn randomly over {*x*_1_,…, *x*_5_} and {+, −, *} respectively – one variable was necessary, since the result of the previous step was considered as the value that always instantiated the first operand in the subexpression. If a variable *x*_1_,…, *x*_5_ or operator {+, −, *} used in an arithmetic expression appeared in a step of the trial, they were encoded by the “high” spiking activity of dedicated 5 neurons (200 Hz), otherwise by their “low’” activity (1 Hz). The numeric inputs to the network were analog values (numbers from [−1, 1] ∈ IR) and encoded by a subpopulation of 250 input spiking neurons. In each step, different numeric values that instantiated the variables *x*_1_,…, *x*_5_ were given, but the network had to learn the binding (“variable - concrete number”), and also to ignore all unnecessary values if the variable did not appear in the current expression. No special modules that would help the binding of variables with numeric values were included. Each analog value used in arithmetic expressions was encoded using a population coding scheme with a population of 50 input neurons. Neurons in this population were tuned to numbers using Gaussian tuning curves where neuron *i* responded maximally with a firing rate of *f*_max_ = 200 Hz if the represented number was equal to its preferred value *m*_*i*_. The preferred values *m*_*i*_ for these 50 neurons were chosen equally spaced in the input domain [−1, 1]. The firing rate *f*_*i*_ of neuron *i* was given by

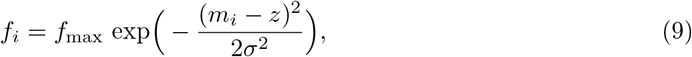

where *z* is the represented value and the standard deviation of *σ* = 0.08 was equal for all input neurons. Each step lasted 50 ms, and consequently, a duration of a trial was 500 ms. From 300 recurrently connected LIF neurons, a half (150 neurons) had an SFA mechanism. Adaptation time constant for each of the neurons with SFA was chosen randomly from a uniform distribution [1, 500]. The output was a linear readout – in each step, output weights were multiplied with the average firing activity of the recurrently connected neurons. This output per step was a single real value.

The network was trained by minimizing the Mean-Squared-Error (MSE) between the network outputs and targets for each step. The final test MSE of 0.0341, and mean absolute error (MAE) of 0.1307 were achieved, calculated for all computational steps in all of, in total, 5000 trials. All test trials had novel sequences of computational steps, that is, test trials were different from those used for training.

## Acknowledgements

**General:** We would like to thank Anand Subramoney for his help and numerous discussions about the experiments done during the master’s thesis of Ceca Kraišniković. These experiments were partly the motivation for the new experiments presented here.

**Funding:** This research was partially supported by the Human Brain Project (Grant Agreement number 785907 and 945539), the SYNCH project (Grant Agreement number 824162) of the European Union, and under partial support by the Austrian Science Fund (FWF) within the ERA-NET CHIST-ERA programme (project SMALL, project number I 4670-N).

